# Joint analysis of matched tumor samples with varying tumor contents improves somatic variant calling in the absence of a germline sample

**DOI:** 10.1101/364943

**Authors:** Rebecca F. Halperin, Winnie S. Liang, Sidharth Kulkarni, Erica E. Tassone, Jonathan Adkins, Daniel Enriquez, Nhan L. Tran, Nicole C. Hank, James Newell, Chinnappa Kodira, Ronald Korn, Michael E. Berens, Seungchan Kim, Sara A. Byron

## Abstract

Archival tumor samples represent a potential rich resource of annotated specimens for translational genomics research. However, standard variant calling approaches require a matched normal sample from the same individual, which is often not available in the retrospective setting, making it difficult to distinguish between true somatic variants and germline variants that are private to the individual. Archival sections often contain adjacent normal tissue, but this normal tissue can include infiltrating tumor cells. Comparative somatic variant callers are designed to exclude variants present in the normal sample, so a novel approach is required to leverage sequencing of adjacent normal tissue for somatic variant calling. Here we present LumosVar 2.0, a software package designed to jointly analyze multiple samples from the same patient. The approach is based on the concept that the allelic fraction of somatic variants, but not germline variants, would be reduced in samples with low tumor content. LumosVar 2.0 estimates allele specific copy number and tumor sample fractions from the data, and uses the model to determine expected allelic fractions for somatic and germline variants and classify variants accordingly. To evaluate using LumosVar 2.0 to jointly call somatic variants with tumor and adjacent normal samples, we used a glioblastoma dataset with matched high tumor content, low tumor content, and germline exome sequencing data (to define true somatic variants) available for each patient. We show that both sensitivity and positive predictive value are improved by analyzing the high tumor and low tumor samples jointly compared to analyzing the samples individually or compared to *in-silico* pooling of the two samples. Finally, we applied this approach to a set of breast and prostate archival tumor samples for which normal samples were not available for germline sequencing, but tumor blocks containing adjacent normal tissue were available for sequencing. Joint analysis using LumosVar 2.0 detected several variants, including known cancer hotspot mutations that were not detected by standard somatic variant calling tools using the adjacent normal as a reference. Together, these results demonstrate the potential utility of leveraging paired tissue samples to improve somatic variant calling when a constitutional DNA sample is not available.

## 1 Introduction

Somatic mutations often drive cancer initiation and progression (Hanahan and Weinberg, 2000). The identification of somatic mutations through next generation sequencing has enabled the identification of cancer driver events in individual patient tumor samples (Allen et al., 2014; Cheng et al., 2015; Frampton et al., 2013). There is also ongoing effort to discover new cancer drivers mutations, particularly in non-coding regions (Khurana et al., 2016). Although sequencing of tumor-associated cancer gene panels and exomes is starting to be adopted in clinical practice to personalize therapy, there is much to learn about how mutation status correlates with response to therapy. Clinically annotated archival tissue collections represent a rich resource for identifying new driver mutations and clarifying how genomic features relate to clinical outcomes (Marrone et al., 2015; Waldron et al., 2012). However, most archival collections do not contain blood samples, or other normal tissue samples from locations distant to the tumor, to use as a constitutional reference. Without a normal tissue sample for comparison, it is difficult to determine which variants are somatic and which are germline (Jones et al., 2015). Innovative approaches are needed to identify somatic variants when normal tissue is not available.

Often, histologically normal tissue is available alongside the tumor biopsy or resection. For instance, surgeons typically remove a margin of adjacent normal tissue when resecting a tumor. This normal tissue can be leveraged for DNA sequencing to identify germline variants. However, it is difficult to know if the adjacent normal tissue is truly free of infiltrating tumor cells. Contamination of the adjacent normal tissue with the tumor tissue during processing could also confound interpretation of the results (Wei et al., 2016). Also, even without infiltrating tumor cells, the adjacent tissue may contain somatic mutations. Field cancerization, where molecular alterations are observed in tissue adjacent to the overt cancer, is thought to be an important risk factor for multifocal and recurrent disease (Dotto, 2014). This phenomenon has been observed in many cancer types including breast (Heaphy et al., 2009) and prostate (Nonn et al., 2009). Even healthy individuals have somatic mutations in normal tissues, and the mutation patterns tend to be similar to those of the cancers arising from that tissue type (Hoang et al., 2016). There even appears to be positive selection for cancer driver mutations in normal skin (Martincorena et al., 2015). Therefore, it is important to consider potential sources of somatic variant contamination when normal tumor-adjacent tissue is used to identify tumor specific somatic variants.

When tumor-only sequencing data is available, researchers have developed various analytic strategies to distinguish germline and somatic variants. One obvious first step to identify somatic variants in tumor-only sequencing data is to filter out the germline variants found in population databases. Jones et al. showed that filtering alone is not sufficient, as each individual typically has an average of 249 private germline variants not found in the population databases that would be incorrectly classified as somatic in tumor-only sequencing (Jones et al., 2015). The number of private germline variants will vary based on the individuals’ ancestry. The private variant rate in a population depends both on how well represented the population is in large scale sequencing projects, as well has the extent to which the population has undergone a recent expansion adding to the diversity of variants (Halperin et al., 2017). More recently, Kalatskaya et al. published a machine-learning approach (ISOWN) to classify somatic and germline variants from tumor-only sequencing data (Kalatskaya et al., 2017). Their approach requires a large training set, and performs best when the training and test datasets are from the same cancer type and patient cohort. In the case of rare cancer types and N of one studies, obtaining such training sets may not be practical. The variant allele fraction, which is the fraction of reads supporting the mutated allele at a given locus, can also help to distinguish somatic from germline variants in impure tumors; the somatic variants should only be present in the tumor cells, leading to a low variant allele fraction, while the germline variants would be present in both the tumor and normal cells in the sample, leading to a variant allele fraction close to 0.5 for heterozygous variants. However, copy number alterations in the tumor can shift both the somatic and germline variant allele fractions, which can lead to considerable overlap in the expected somatic and germline variant allele fractions and greatly reduce the power to detect somatic variants (Figure 1). Thus, there is a need for new bioinformatics methods to call germline and somatic variants from tumor samples with high sensitivity and precision, even in the absence of a germline sample.

**Figure 1.**
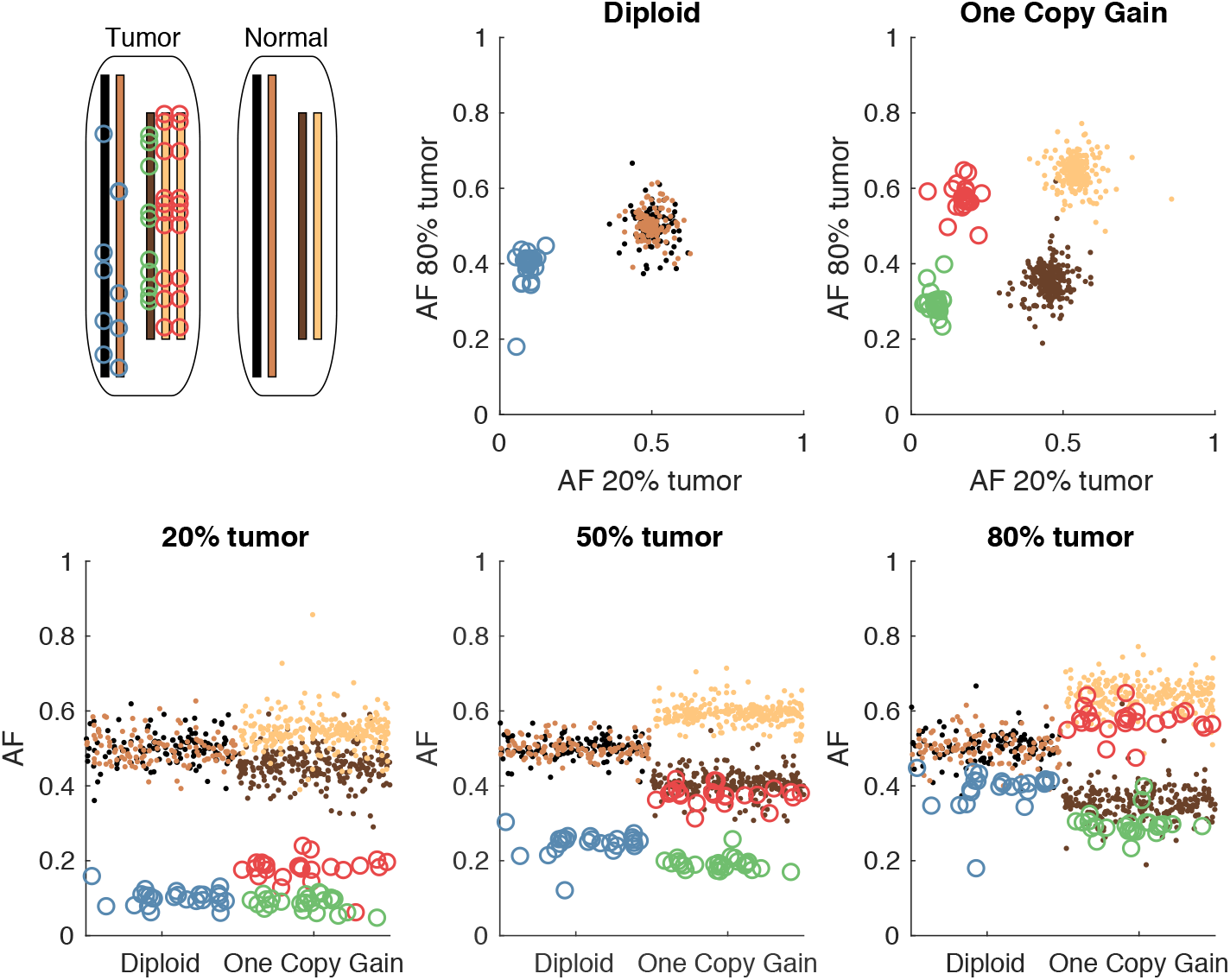
Somatic and germline variant allelic fractions example. A) Two chromosomes are illustrated for this example. Both chromosomes are present in the diploid state in the normal cell. In the tumor cell, one chromosome is in the diploid state, and the other shows one-copy gain. Blue circles represent somatic variants on the diploid chromosome, green and red circles represent somatic variants on the minor and major alleles of the gained chromosome, respectively. Simulated allelic fractions of germline variants (brown/tan) and somatic variants are plotted for a simulated 20% tumor (D), 50% tumor (E) and 80% tumor (F) by chromosome position. In the 50% tumor example, somatic variants could easily be distinguished from germline on the diploid chromosome, but on the copy number gain chromosome, the allelic fractions of the somatic variants on the major allele overlap with the germline variants. By using both the 20% and 80% tumor samples, the somatic variants can be separated from the germline variants by allelic fraction on both the diploid chromosome (B) and the copy number gain chromosome (C).

**Figure 2.**
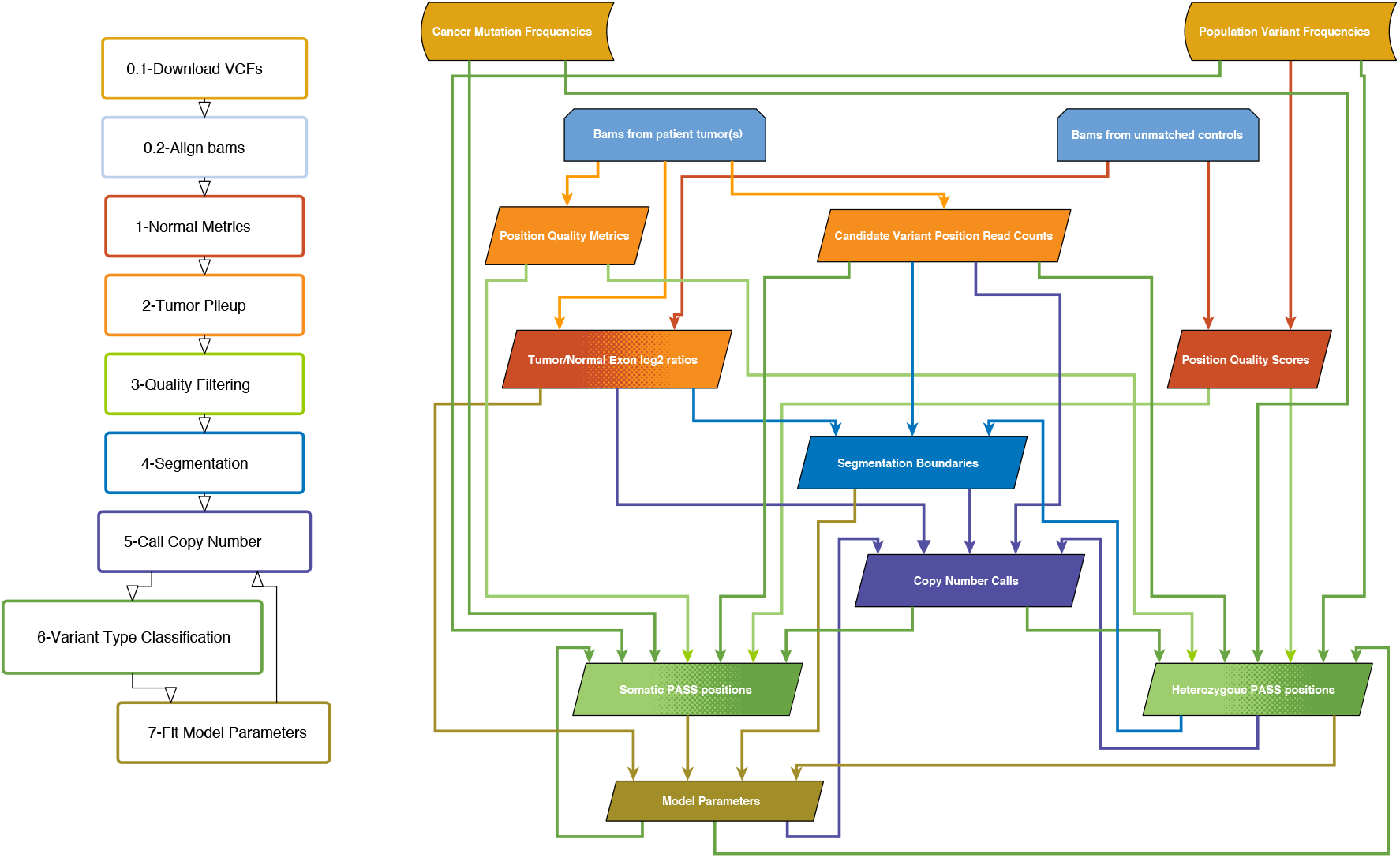
Overview of LumosVar 2.0 Analysis. The flow-chart on the left show the main steps in the analysis. Steps 0.1 and 0.2 are data preparation, and steps 1-7 are performed by lumosVar 2.0. The graph on the right illustrates the main inputs and outputs of each step. The color of the arrows coming from each box indicates the steps where that data is used as input, and the color of each box indicates the step where the data is generated.

Here, we present a new bioinformatics approach (LumosVar 2.0) that leverages adjacent normal tissue from tumor biopsies and permits somatic mutations to be present in the adjacent tissue. Similar to our previous approach (LumosVar – single sample-based variant caller) (Halperin et al., 2017), we model allelic copy number to determine the expected allelic fractions for somatic and germline variants as well as incorporate population database frequencies to call variants as somatic or germline. We have extended the approach to find the joint probability of somatic and germline mutations across multiple samples from the same patient. We hypothesize that the patterns of allelic fractions across samples of different purities will be more informative than any individual sample in distinguishing somatic from germline variants (Figure 1). To test this hypothesis, we compare two approaches 1) jointly calling variants using two samples of different purities (joint approach) and 2) pooling the two samples resulting in one sample with twice the sequencing depth and the average of the purities (pooled approach). First, we used simulations to systematically evaluate the effects of tumor purity and copy number states for the two approaches. Next, we looked at a set of glioblastoma (GBM) patient samples where we have sequencing data for contrast-enhancing region (CE, high fraction of tumor cells) and non-enhancing region (NE, low fraction of tumor cells) biopsies, as well as sequencing data from peripheral blood samples to establish true somatic calls. Finally, we applied our method to an archival cohort of breast and prostate samples where FFPE sections from the tumor biopsies or resections were the only tissues available.

## 2 Methods

### 2.1 Simulations

We used simulated data to systematically determine how the purity of the two tumor samples, the copy number, and the read depth affect our ability to detect somatic variants. The simulations were performed as previously described (Halperin et al., 2017), where the total read depths were drawn from a log normal distribution and the number of reads supporting the variant were drawn from a binomial distribution. To evaluate how the joint calling approach compares to the single sample approach, we added the read depths of each pair of simulated tumor samples used jointly.

### 2.2 Evaluation dataset

A set of previously collected and de-identified whole exome data from seven recurrent GBM patients was used to evaluate the approach. Each patient dataset contained exome sequencing data for CE biopsies (high tumor content), NE biopsies (low tumor content), and peripheral blood (germline). The acquisition and sequencing of these samples was previously described (Byron et al., 2017). The consensus of three comparative somatic variant callers (seurat (Christoforides et al., 2013), strelka (Saunders et al., 2012a), and mutect (Cibulskis et al., 2013)) was used to define the true somatic variants. We determined the number of likely germline false positives based on the number of heterozygous variants called by HaplotypeCaller (DePristo et al., 2011) that were not found in dbSNP (Sherry et al., 2001).

### 2.3 Application to archival sample sets

De-identified FFPE tissue sections, clinical data, and pathology data were acquired for twenty breast cancer patients and twenty prostate cancer patients from HonorHealth Scottsdale Shea Medical Center, in accordance with local institutional review boards and in compliance with the Health Insurance Portability and Accountability Act (HIPAA).

IRB approval included a waiver of informed consent for the prostate cancer patients. Patients in the breast cancer cohort provided signed consent to participate in the study. Retrospective analysis was performed using archival samples from treatment-naïve, invasive breast carcinomas or treatment-naïve prostate adenocarcinomas.

Breast tumors were collected following routine clinical lumpectomy or mastectomy, from women diagnosed with ER-positive, invasive mammary carcinoma between 2010 and 2016 at HonorHealth Scottsdale. Median age of diagnosis was 65 years and ranged from 39 to 86 years. All tumors were classified by pathology as estrogen receptor-positive. Nineteen of the twenty tumors were classified as HER2-negative. The breast tumor cohort spanned AJCC stages (IA-IV). Prostate tumors were collected following radical prostatectomy for men diagnosed with pancreatic adenocarcinoma between 2012 and 2016 at HonorHealth Scottsdale. Median age of diagnosis was 67 years, ranging from 57 to 74 years. Eighteen of the twenty tumors had a Gleason score of seven or greater. ER/PR/HER2 status (breast tumors), Gleason score (prostate tumors), histological type, tumor stage, treatment history, and clinical outcome, including progression-free survival and overall survival, were collected from medical records and the de-identified data was provided for this study. Pathology review (J.N.) identified a tissue block with high tumor content and a tissue block with a region considered to have low tumor content for each patient. Five 10-micron sections were provided for each sample (5 high tumor content; 5 low tumor content). The Qiagen GeneRead FFPE DNA Kit (cat# 180134) was used to isolate DNA from FFPE breast and prostate cancer tumor specimens (N=80) following the manufacturer’s protocol.

Exome libraries were constructed from 200ng of DNA (DIN=3-5) using KAPA Biosystems’ Hyper Prep Kit (cat#KK8504) and Agilent’s SureSelectXT V5 baits, containing custom content, following the manufacturer’s protocols. Custom bait content included copy number probes distributed across the entire genome, along with additional probes targeting tumor suppressor genes and genes involved in common cancer translocations to enable structural analysis. Libraries were equimolarly pooled, quantitated, and sequenced by synthesis on the Illumina HiSeq 4000 for paired 82bp reads.

### 2.4 Variant caller overview

We previously created a single-sample strategy (lumosVar) to call somatic variants in impure tumor samples based on the differences in allelic frequency between the somatic and germline variants (Halperin et al., 2017). Here, we describe an extension of lumosVar to jointly analyze multiple samples from the same patient. The lumosVar 2.0 analysis has seven main steps. First, a set of unmatched control samples is analyzed for position quality scores and average read depth, as previously described (Halperin et al., 2017). Second, read counts and quality metrics are extracted from the tumor bams. Third, quality scores are calculated for each candidate variant position. Fourth, segmentation is performed to define regions that have similar tumor/normal read depth ratios and B-allele fractions. The fifth step involves finding the most likely allele-specific copy number state for each segment. The sixth step involves classifying each candidate variant position as somatic, germline heterozygous, or homozygous. The final step entails optimization of the model parameters. The caller iterates between steps five, six, and seven until the solution converges.

### 2.5 Quality Classification and Filtering

We used sixteen quality metrics in a quadratic discriminant model to determine the posterior probability that each position belongs to a PASS group, as was previously used in LumosVar. The same quality metrics and thresholds are used in LumosVar 2.0 as were used in LumosVar to assign candidate variant positions to PASS and REJECT training groups (Halperin et al., 2017). As previously, here we also fit the model separately on candidate indel and point mutation positions. Since the quality metrics for the B allele are not relevant for homozygous positions, after the joint variant calling is performed as described below, we repeat the quality classification step fitting homozygous positions separately after setting all the quality metrics to only use the “A” allele and setting the difference metrics to zero. The model is fit independently on each sample, and then we calculate a trust score by taking the geometric mean of the posterior probability of belonging to the PASS group weighted by the number of reads supporting the variant across the samples. Candidate positions with a trust score greater than a threshold (T_pass_) are considered for classification as somatic or germline variants as described in the joint somatic variant calling section below.

### 2.6 Segmentation

Prior to fitting the copy number model, segmentation is performed. We use the circular binary segmentation algorithm implemented in the Matlab Bioinformatics toolbox to segment both the tumor to normal read depth log2 ratio of each exon and the B-allele frequencies of common heterozygous variants. Segmentation is performed independently on each sample. We combine all of the segmentation boundaries from all of the samples for both the read depth log2 ratio and B-allele frequency segmentation, and then remove non-significant segments as follows. A two-sample t-test is used to compare each pair of adjacent segments for both the read depth log2 ratios and the B-allele fractions for each sample. For each segmentation boundary, the geometric mean is used to combine the p-values across the samples and data types. The segmentation boundary with the highest geometric mean p-value is removed, and the t-tests are then performed on the newly merged segment with its neighbors. This process is continued until of the segmentation boundaries have geometric mean p-values less than the segmentation significance threshold (*α* _*seg*_).

### 2.7 Expectation Maximization

We use an expectation maximization approach to fit the model parameters and call variants. In the initial iteration, heuristics are used to find reasonable values of the model parameters. In the expectation step, the model parameters are used to identify somatic and germline heterozygous variant positions. Identifying these variant positions involves finding the copy number states of each segment, joint variant type classification, and variant quality filtering, all as described below. Using those variant positions, values of the model parameters that maximize the likelihood of the data are found.

### 2.8 Initial Parameter Values

The parameter f is a matrix with the number of rows corresponding to the number of clonal variant groups (K) and the number of columns corresponding the number of samples (J). The initial value of f is set such that there is a main clonal variant group that has a high sample fraction in all samples, there are J clonal variant group that clonal in each sample and low in the other samples, and there are another J clonal variant groups that are sub-clonal in each sample and very low in the other samples. The centering and spread parameters (C and W) are both vectors of length J, and their initial values are determined as previously described (Halperin et al., 2017).

### 2.9 Copy number state assignments

The copy number state of each (g) may be described by the total copy number (N_g_), the minor allele copy number (M_g_), and the index of clonal variant group of the segment (k_g_). The values of N_g_, M_g_, and k_g_ are found that maximize a sum of log likelihoods for the segment (SLL_g_).

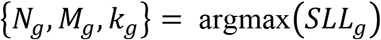

This sum includes the likelihoods of the exon mean read counts (L_xj_), heterozygous variant read counts (L_yj_), number of common germline variant positions that would be called germline heterozygous (*L_Ydg_*) or somatic (*L_Ydg_*), as well as the prior probabilities of the copy number states (π (N), π (M)) and sample fraction difference 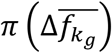

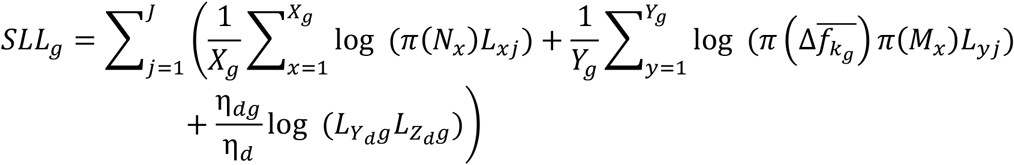

The likelihood calculations are defined below.

### 2.10 Parameter fitting procedure

The values of f, W, and C are found that maximize the sum of segment log likelihoods (SLL_g_) and somatic variant log-likelihoods (L_z_). Since the number of clonal variant groups (K) changes the degrees of freedom of the model, as f is a J by K matrix, we include a penalty term for increasing K.

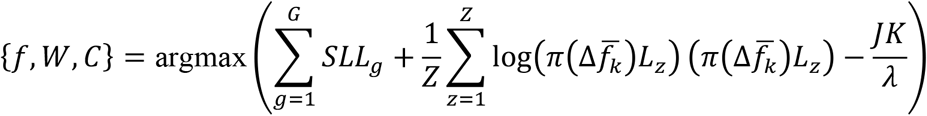

In order to more efficiently search the parameter space, we use the parameter values from the previous EM iteration (or the initial values in the first iteration) as the starting point for the parameter optimization. Since the heuristic used to find the initial value of C may be incorrect, particularly for higher ploidy genomes, other values of the centering parameter are also tested, and the best one is used as a starting point for optimization of all parameters. In order to find a reasonable starting point for adding an additional clonal variant group, we use the previous f matrix and test a set of random values for the additional column. We use the best one as the starting point for optimizing all of the parameters. If the maximum likelihood score improves, the procedure is repeated with an additional clonal variant group. This process continues until adding a clonal variant group fails to improve the likelihood score. Since an adding a clonal variant group may make a previous clonal variant group less important to the model, we also test removing each clonal variant group, and then do another round of optimization of all parameters. If this results in a new maximum, then the procedure will be repeated removing another clonal variant group. Once removing clonal variant groups no longer improves the model, the procedure returns to re-centering. The re-centering, adding clonal variant groups, and removing clonal variant groups is repeated until there are ξ consecutive iterations with no new maximum found.

### 2.11 Likelihood calculations

As in lumosVar, the likelihood of the exon mean read depths are modeled as a Poisson distribution, and the somatic and germline heterozygous read counts are modeled as beta binomial distributions.

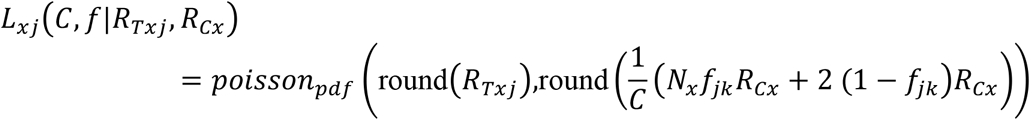

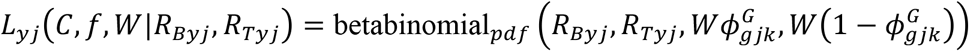

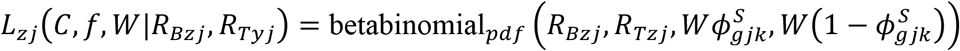

The expected germline heterozygous variant allele fraction is determined as follows.

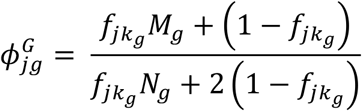

LumosVar 2.0 considers three possible scenarios when finding the expected somatic variant allele fraction: 1) variant is in the same clonal variant group as the copy number alteration effecting the segment and is on the minor allele (*k_zj_* ≡ *k_gj_* Λ *A_z_* ≡ 1), 2) variant is in the same clonal variant group as the copy number alteration effecting the segment and is on the major allele (*k_zj_* ≡ *k_gj_* Λ *A_z_* ≡ 2), or 3) variant is in a non copy number altered clonal variant (*k_zj_ ≠ k_gi_*).

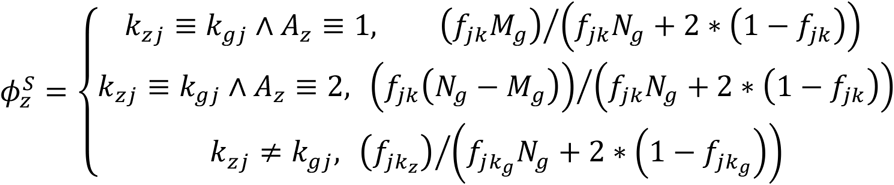

The maximum likelihood is used to determine the clonal variant group assignment and allele of each somatic variant.

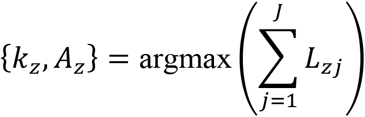

The probability of detecting a heterozygous variant in each segment is calculated based on the cumulative probability of observing at least the minimum number of reads required to be considered a candidate variant position (R_B-min_), given the mean read depth in the segment (R_T_) and the expected allele fraction of a heterozygous variant in that segment 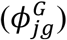.

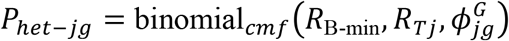

In order to determine if parameter values would result in reasonable variant counts, the variant type classification is performed at common germline variant positions. The likelihood of detecting fewer than the observed number of heterozygous variants in a segment (Y_dg_) is modeled as the cumulative probability from a binomial distribution with Y_dg_ successes, the number of bases examined in the segment (η_dg_), and *p_het_* probability of success.

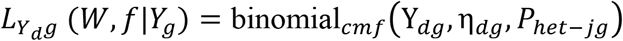

In order to penalize models that would result in germline variants being called somatic, we then determine the likelihood of finding that many or more somatic variants in germline variant positions based on the cumulative probability from a binomial distribution with Z_dg_ somatic variants detected of η_*dg*_ database variant positions tested, with a probability of success of *ρ_SNV_*.

*L_Zdg_ (C,W,f|Z_dg_)* = binomial*_cmf_* (Z*_dg,_η_dg,_ρ_SNV_* In lumosVar, we set a prior distribution on f in order to favor models where the sample fractions are close to what is expected. In lumosVar 2.0, we set a prior distribution on the difference in f across the samples to favor models where the sample fractions differ as much or more than the expected sample fractions. The mean difference is found for prior tumor sample fractions (*f π*) as follows:

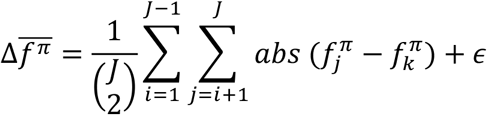

The mean difference of the sample fractions for each clone is found as similarly.

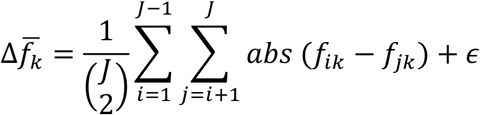

The prior probability that the sample fractions for each clone have a mean difference as much as or greater than observed is calculated from a beta distribution with a mode of the difference in the prior tumor sample fractions.

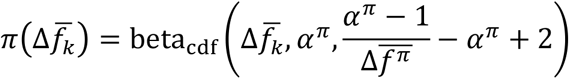

### 2.12 Joint Variant Type Classification

The probability of observing the read counts in each sample (k) given that the variant is somatic (P (D_k_|S)), germline heterozygous (P (D_k_ |G_AB_)), germline homozygous (P (D_k_ |G_AA_)), or another genotype (P (D_k_ |O)) are calculated as previously described. The prior probabilities are also calculated as previously described (Halperin et al., 2017). The product of the conditional probabilities across the set of samples gives the joint probability of each variant type, given all the samples’ data, as we assume that the read counts for each sample are independent. The posterior probability that a position has a somatic variant given all the samples’ data is calculated as shown below.

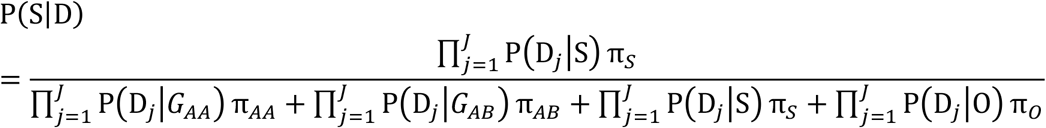

### 2.12.1 Implementation and availability

A custom pileup engine was written in C using htslib (https://github.com/tgen/gvm). The pileup engine extracts the mean exon read depths and calculates the quality scores form the unmatched control bams, as well as extracts the read counts and quality metrics from the tumor bams. The rest of the lumosVar 2.0 analysis was written in Matlab (https://github.com/tgen/lumosVar2). A precompiled binary is provided which enables users to run lumosVar 2.0 without a Matlab license.

## 3 Results

### 3.1 Simulations: comparison between pooled and joint approaches

Simulations were performed to determine how the tumor purity and copy number states affect the power to detect somatic variants in the joint approach, and how the power compares to the pooled approach. As previously shown, the pooled single sample approach performs best with a sample of intermediate tumor purity for variants in diploid regions, but copy number variation leads to situations where the expected somatic and germline allele fractions are very similar, making it difficult to classify somatic variants using a single sample (Halperin et al., 2017). From the simulation results, we can see that the joint approach mitigates this limitation, and only provides poor detection when both samples fall into a range where the expected somatic and germline allele fractions are very similar (Figure 3). The joint approach generally only requires low-to-moderate coverage when one sample has low tumor content and the other sample has moderate-to-high tumor content.

**Figure 3.**
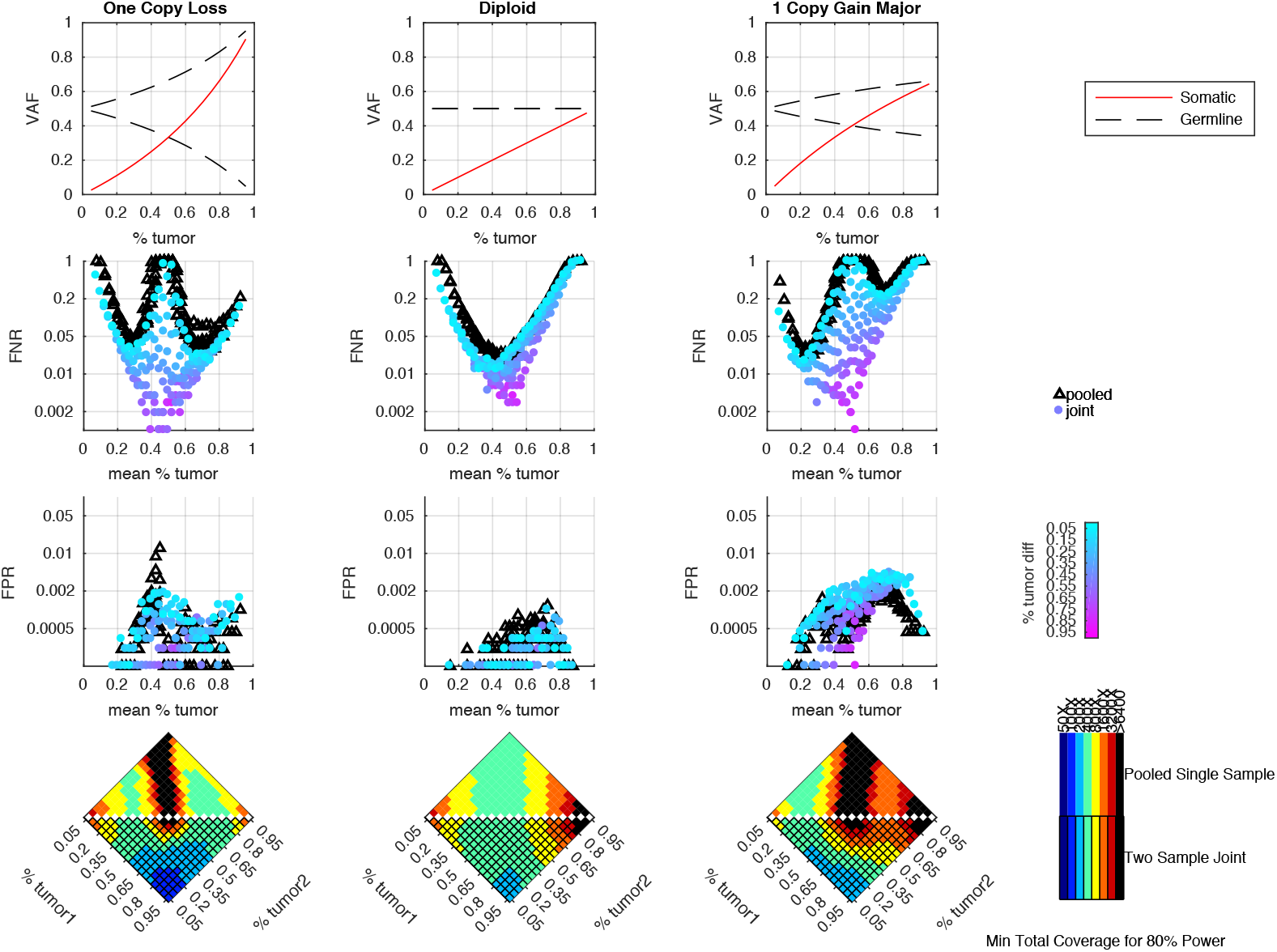
Simulation results comparing pooled and joint approaches. Top row of graphs shows the expected allele frequency of somatic (red) and germline variants (black) by tumor content (x-axis) for different copy number states. The middle two rows of graphs are based on simulation results using a mean coverage of 200X per sample (400X pooled). They show the false negative rate (FNR – simulated somatic variants not called somatic) and false positive rate (FPR – simulated germline heterozygous variants falsely called somatic) plotted by mean tumor content for the pooled (black triangles) and joint (colored circles) approaches. For the joint approach, the color of the circles represents the difference in tumor content between the two samples analyzed jointly. The bottom set of graphs shows the coverage required to detect at least 80% of the simulated somatic variants using two samples of different tumor content (shown on the x and y axis) using a joint approach (lower triangle of each heatmap) or using a single-sample approach on a merged sample with a tumor content that is the average of the two samples and coverage that is the sum of the two samples (upper triangle of heatmap). The color indicates the mean target coverage in the pooled approach, or the sum of the mean target coverage in the two-sample joint approach. Black squares indicate that less than 80% of the somatic variants were detected at the highest coverage simulated (6400X).

### 3.2 Clonal variant groups

While the single sample version of lumosVar also assigned somatic mutations and copy number alterations to clonal variant groups, these clonal variant groups become much more informative when looking at more than one patient sample. An example patient’s results are shown in Figure 4. There are three clonal variant groups found in this patient, one that appears clonal in both samples (blue), one that appears sub-clonal in the enhancing biopsy and not detected in the non-enhancing biopsy (red), and one that appears clonal in the non-enhancing biopsy and sub-clonal in the enhancing biopsy (green). From these groups we can infer that the enhancing sample likely has two clones, one with the blue and red variants, and a second with the blue and green variants, while the non-enhancing sample appears to only have the clone with the blue and green variants. This patient also illustrates why the joint calling approach is advantageous to detect somatic variants if the germline was not available. With only the enhancing sample, the blue variants would be difficult to differentiate from the germline variants. If the non-enhancing sample were used as a reference in standard paired somatic variant calling, only the red variants would likely be detected.

**Figure 4.**
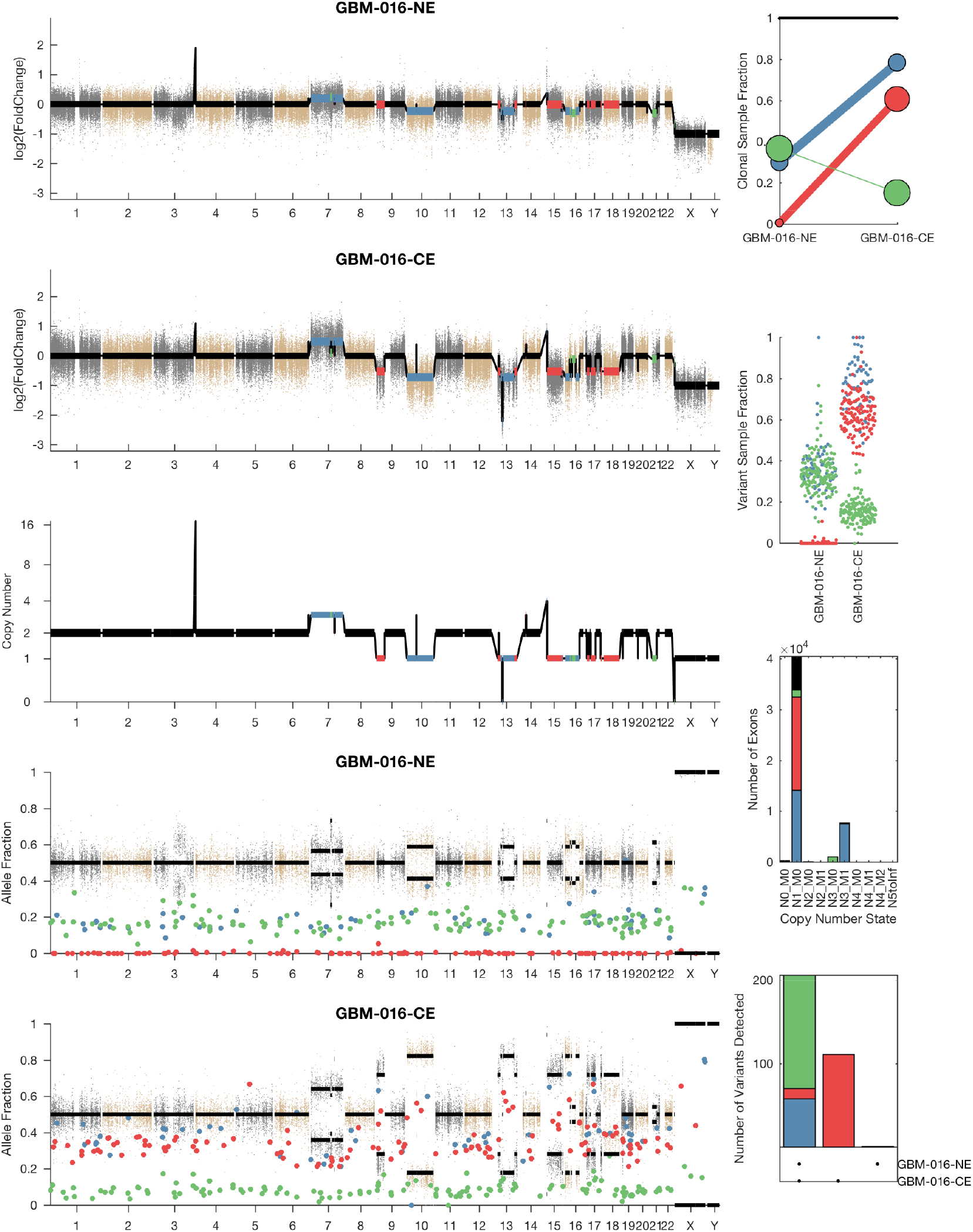
Example LumosVar 2.0 Output. A) Log2 fold change of the mean exon read depths compared to the unmatched controls. B) The estimated integer copy number states are plotted for each genomic segment by chromosome position. C) The variant allele fractions are plotted by chromosome position. The grey and brown dots represent variants called as germline heterozygous by lumosVar 2.0 and the large colored dots represent variants called somatic by lumosVar 2.0. D) Summary of the clonal variant group patterns. The thickness of the lines represents the proportion of copy number events assigned to each group and the size of each circle is proportional to the number mutations assigned to each group. E) Sample fraction distribution of somatic mutations. F) Number of exons determined to be in each copy number state. G) Number of somatic mutations detected in both samples (left bar), enhancing only (middle bar), and nonenhancing only (right bar). On all plots, the colors indicate the clonal variant group.

### 3.3 Evaluation: real patients

To evaluate LumosVar 2.0 on real data, recurrent glioblastoma patients that had whole exome sequencing data available for two samples of different tumor contents (from contrast enhancing and non-enhancing biopsies), as well as germline sequencing data (from peripheral blood), were identified. Three variant calling approaches were compared: (1) A filtering approach, where heterozygous germline variants not found in dbSNP were considered false positives, and somatic variants found in dbSNP were considered false negatives, (2) a pooled approach where the data for the high tumor content and low tumor content samples are combined *in-silico*, and (3) joint analysis of the paired high tumor content and low tumor content samples. Both the pooled and joint approaches use the LumosVar 2.0 software for variant calling. We find that the filtering approach consistently has better sensitivity, but much lower precision, and lower F1 scores (harmonic mean of sensitivity and precision) than both the pooled and joint analyses (Table 4). This is consistent with our previous findings that private germline variants result in a high number of false positives using a filtering approach (Halperin et al., 2017). In most of the samples, we find modest improvements in sensitivity, precision, and F1 scores in the joint approach compared to the pooled. From the simulations, we would have expected to see similar precision and more consistent improvements in sensitivity. In order to more carefully evaluate where the joint approach and pooled approach are performing differently in detecting variants, we examined the sample fractions of variants that are true positives, false positives, and false negatives in each approach (Figure 5). We find that the pooled approach has more false positive variants that have similar allelic fractions in the CE and NE biopsies. We hypothesize that these variants have unexpected allelic fractions do to mapping noise or copy number call errors that would not be modeled in the simulations. The joint approach is better at avoiding these calls, as the allelic fractions do not fit the patterns of the clonal variant groups found in the patient. However, the joint approach also misses some true somatic variants that do not fit the patterns of clonal variant groups found in the patient, such as a set of lower sample fraction variants in GBM-003. GBM-014 is the only patient where the pooled approach outperforms the joint approach. This patient also appears to have the smallest difference in tumor content between the two biopsies as well as the most complex copy number profile of this set of patients, both factors that likely contribute to the poor performance.

**Figure 5.**
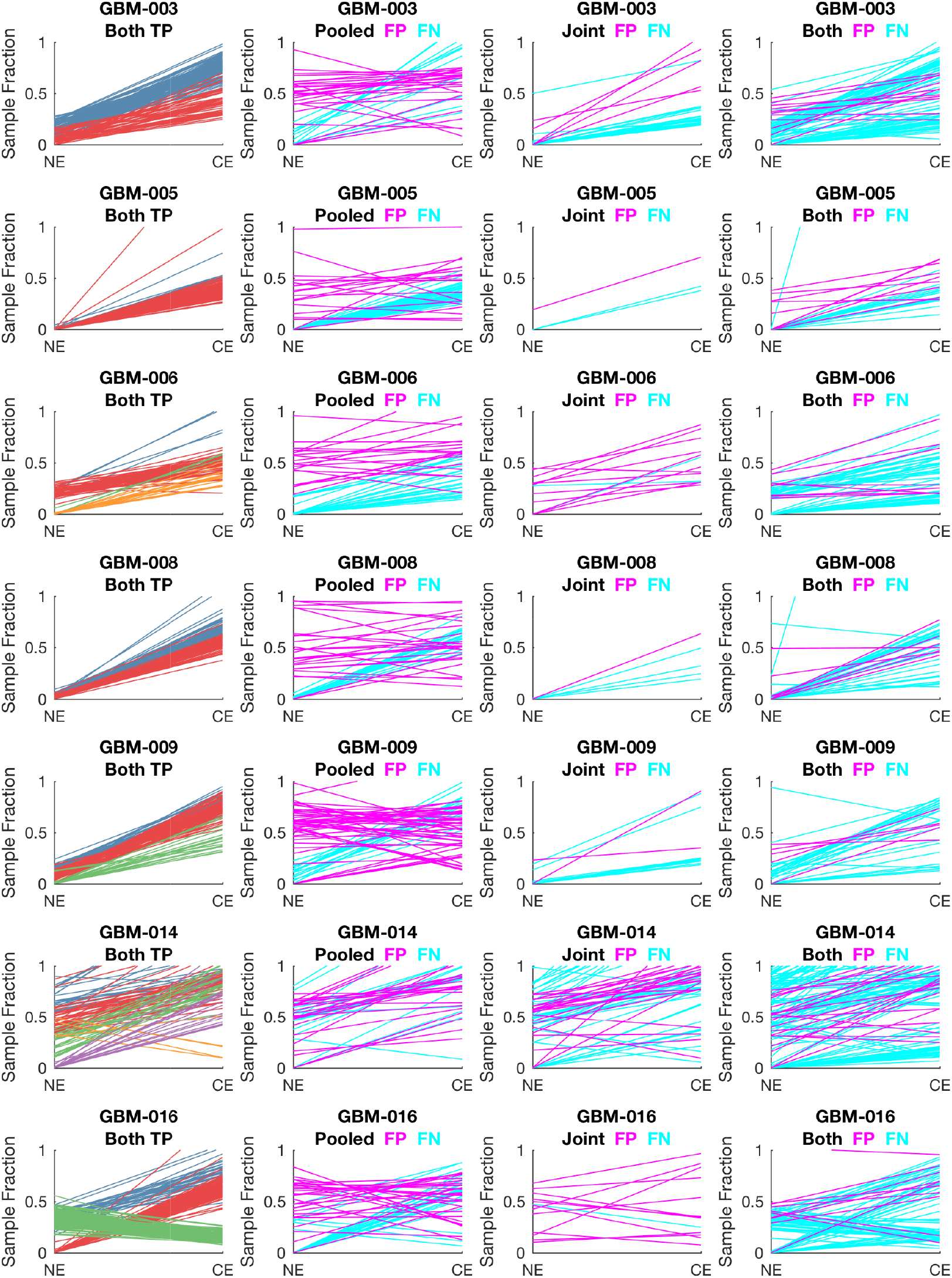
Comparison of Variants Called in Pooled vs. Joint Approach. The first column of graphs shows the estimated sample fractions of true somatic variants that were detected by both the pooled and joint approaches. The variants are colored by clonal variant groups. The other three columns show the sample fractions of variants that were called incorrectly only in the pooled approach (column 2), only in the joint approach (column 3), or incorrectly in both approaches (column 4). False positives variants are shown in magenta and false negatives in cyan.

**Table 1.** Patient Characteristics.

| Patient Id | Cancer Type | Low Tumor Source<sup>a</sup> | Mean Target Coverage |  |  |
| --- | --- | --- | --- | --- | --- |
|  |  |  | Low Tumor | High Tumor | Peripheral Blood |
| GBM-003 | GBM | NEB | 183 | 387 | 177 |
| GBM-005 | GBM | NEB | 482 | 404 | 144 |
| GBM-006 | GBM | NEB | 441 | 248 | 115 |
| GBM-008 | GBM | NEB | 203 | 424 | 186 |
| GBM-009 | GBM | NEB | 199 | 387 | 153 |
| GBM-014 | GBM | NEB | 223 | 366 | 114 |
| GBM-016 | GBM | NEB | 416 | 376 | 261 |
| BHH01 | Breast | ANWS | 255 | 268 | NA |
| BHH02 | Breast | ANMD | 296 | 350 | NA |
| BHH03 | Breast | ANMD | 228 | 302 | NA |
| BHH04 | Breast | ANMD | 269 | 296 | NA |
| BHH06 | Breast | ANMD | 249 | 336 | NA |
| BHH09 | Breast | ANMD | 312 | 290 | NA |
| BHH11 | Breast | ANMD | 249 | 282 | NA |
| BHH15 | Breast | ANWS | 211 | 323 | NA |
| BHH16 | Breast | ANWS | 294 | 289 | NA |
| BHH21 | Breast | ANMD | 286 | 331 | NA |
| BHH22 | Breast | ANWS | 226 | 301 | NA |
| BHH24 | Breast | ANWS | 229 | 297 | NA |
| BHH25 | Breast | ANWS | 278 | 275 | NA |
| BHH26 | Breast | ANMD | 301 | 291 | NA |
| BHH27 | Breast | ANWS | 236 | 328 | NA |
| BHH28 | Breast | ANWS | 275 | 264 | NA |
| BHH08 | Breast | ANWS | 286 | 288 | NA |
| BHH18 | Breast | ANWS | 287 | 261 | NA |
| BHH20 | Breast | ANWS | 185 | 326 | NA |
| BHH23 | Breast | ANWS | 273 | 243 | NA |
| HHP01 | Prostate | ANWS | 202 | 216 | NA |
| HHP02 | Prostate | ANWS | 206 | 221 | NA |
| HHP03 | Prostate | ANWS | 261 | 184 | NA |
| HHP04 | Prostate | ANWS | 312 | 241 | NA |
| HHP05 | Prostate | ANWS | 247 | 232 | NA |
| HHP06 | Prostate | ANWS | 294 | 214 | NA |
| HHP07 | Prostate | ANWS | 265 | 254 | NA |
| HHP08 | Prostate | ANWS | 287 | 247 | NA |
| HHP09 | Prostate | ANWS | 302 | 228 | NA |
| HHP10 | Prostate | ANWS | 328 | 277 | NA |
| HHP11 | Prostate | ANWS | 329 | 274 | NA |
| HHP12 | Prostate | ANWS | 238 | 239 | NA |
| HHP13 | Prostate | ANWS | 299 | 285 | NA |
| HHP14 | Prostate | ANWS | 269 | 269 | NA |
| HHP16 | Prostate | ANWS | 363 | 330 | NA |
| HHP17 | Prostate | ANWS | 255 | 316 | NA |
| HHP18 | Prostate | ANWS | 224 | 258 | NA |
| HHP19 | Prostate | ANWS | 241 | 281 | NA |
| HHP20 | Prostate | ANWS | 208 | 307 | NA |
| HHP21 | Prostate | ANWS | 213 | 263 | NA |

<sup>a</sup>NEB: non-enhancing biopsy, ANWS: adjacent normal whole slide, ANMD: adjacent normal macrodissected.

**Table 2.**
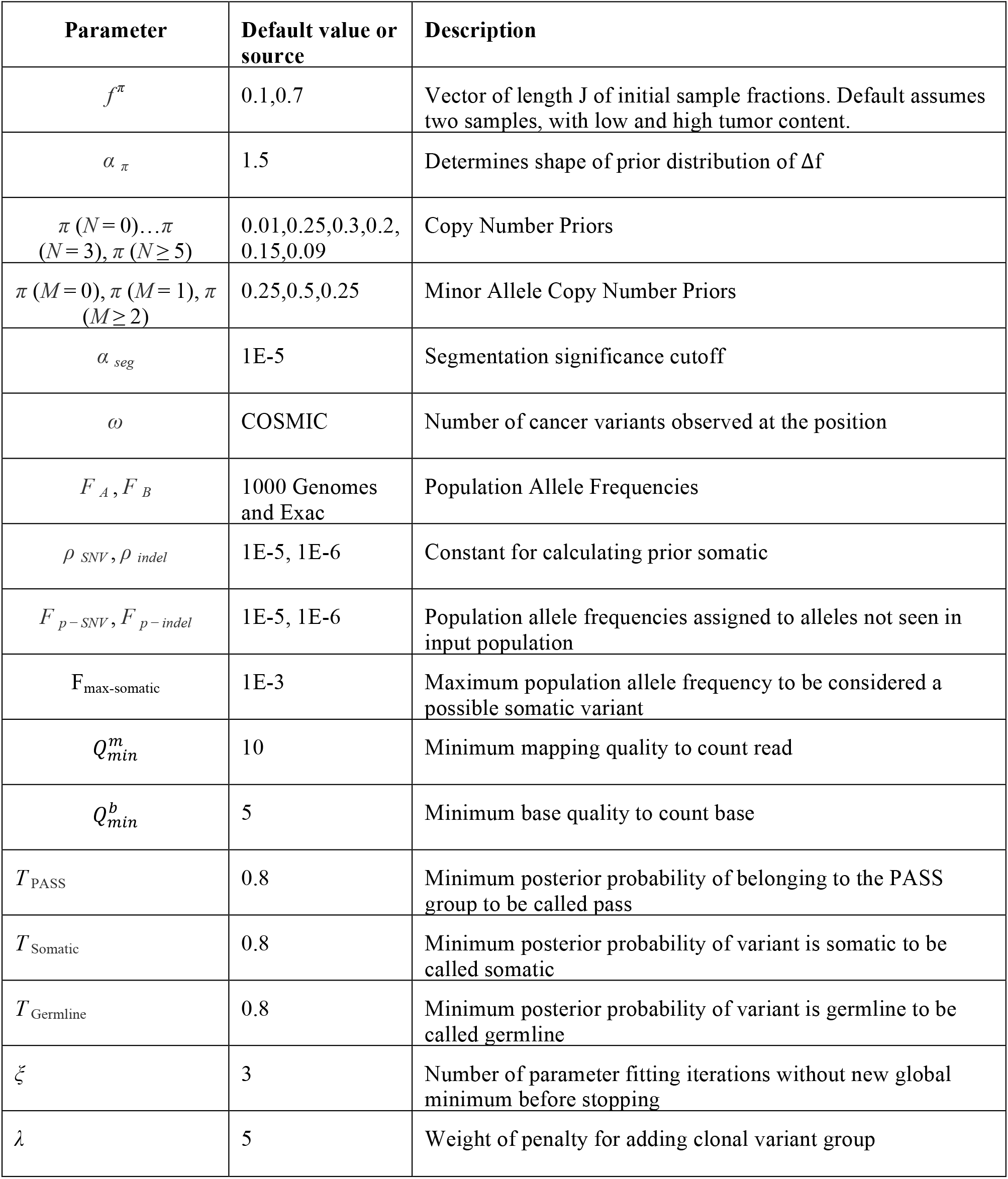
Parameters and Default Values.

**Table 3.**
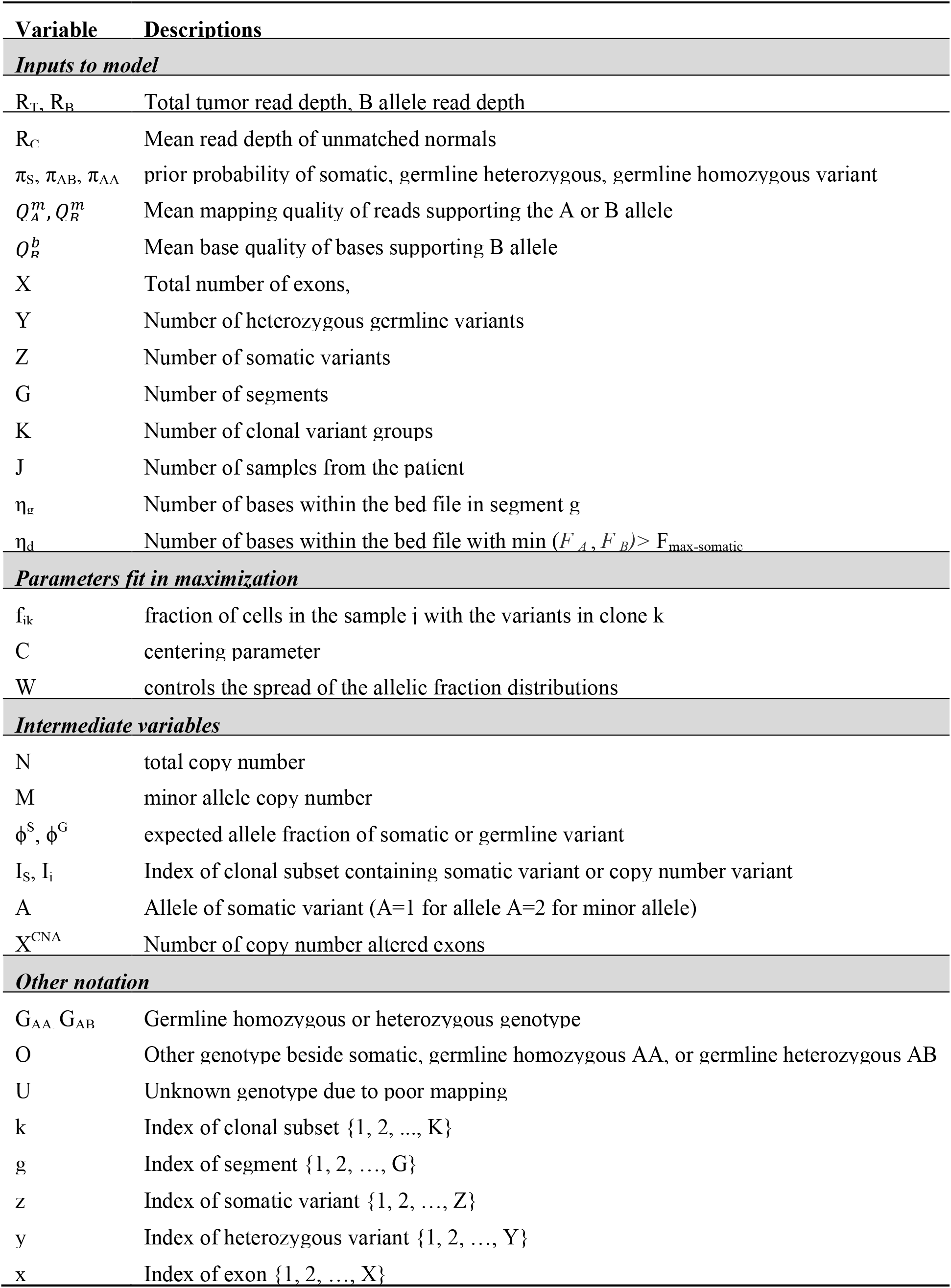
Other Parameters and Notation.

**Table 4.**
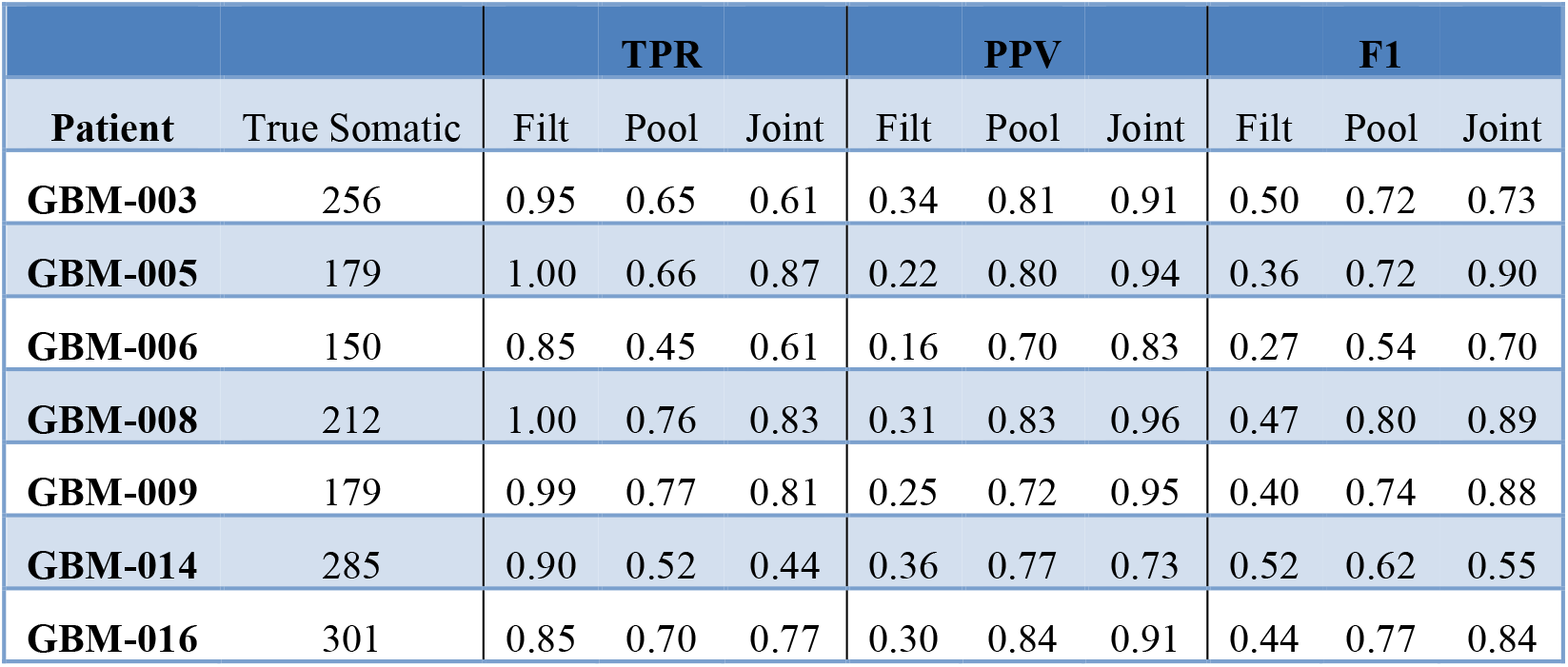
Evaluation Results. The number of true somatic variants found in each patient, as well as the sensitivity (TPR), precision (PPV), and F1 score are shown for the filtering (Filt), single sample (Pool), and lumosVar 2.0 (Joint) approaches.

### 3.4 Application to archival samples

We applied our methods to archival breast cancer and prostate samples, where only FFPE tissue sections from biopsies or surgical tumor resections were available. For eight of the breast cancer patients, whole slides with adjacent normal tissue were not available or did not have sufficient DNA yield, so adjacent normal areas were macro-dissected from tumor-containing slides. For the remaining patients, DNA was isolated from whole slides from additional FFPE blocks containing adjacent normal tissue. For two breast cancer cases (BHH02, BHH27), the additional “low tumor” blocks were from the contralateral breast following double mastectomy, though BHH02 was one of the eight patients that required macro-dissections of the tumor-containing slide to get sufficient DNA for the adjacent normal sequencing. Where macro-dissection was used to obtain the normal tissue samples, most of the somatic variants called were detected the adjacent normal (median of 98%). For the patients where adjacent normal tissue was obtained from separate slides, most patients still had some somatic variants detected in the normal tissue (median 35% - Figure 6).d

**Figure 6.**
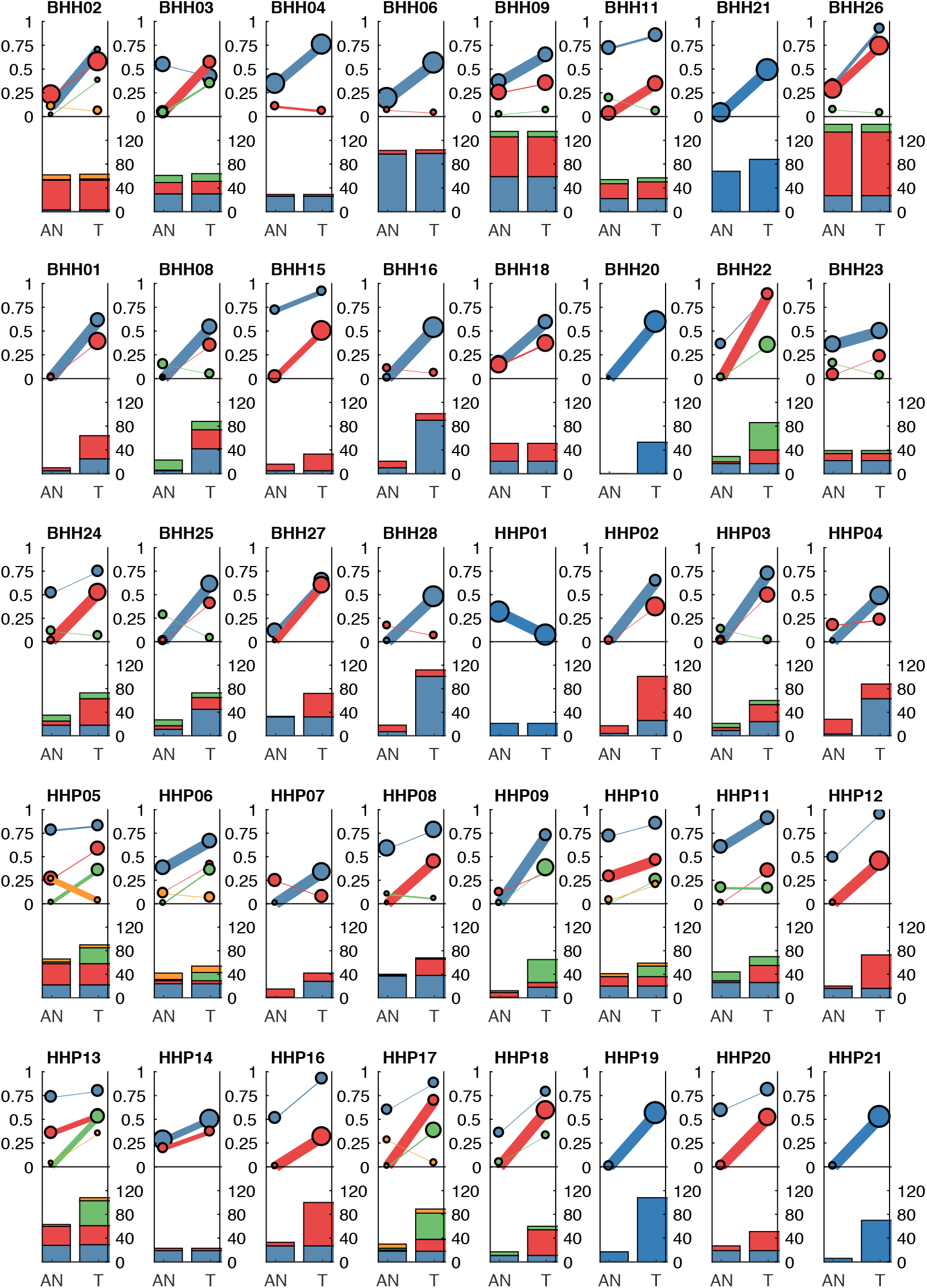
Clonal patterns and variant counts detected by LumosVar 2.0 in the archival dataset. The top half of each plot shows the summary of the clonal variant group patterns for each patient. Each line represents a clonal variant group and the thickness of the lines represents the proportion of copy number events assigned to each group and the size of each circle is proportional to the number mutations assigned to each group. The bottom half of each plot shows the number of somatic variants detected in the adjacent normal (AN) and tumor (T) samples, with the colors corresponding the clonal variant groups. The eight patients in the top row had the adjacent normal tissue macrodissected from tumor containing slides and these patients typically have similar number of variants detected in the tumor and adjacent normal.

**Figure 7.**
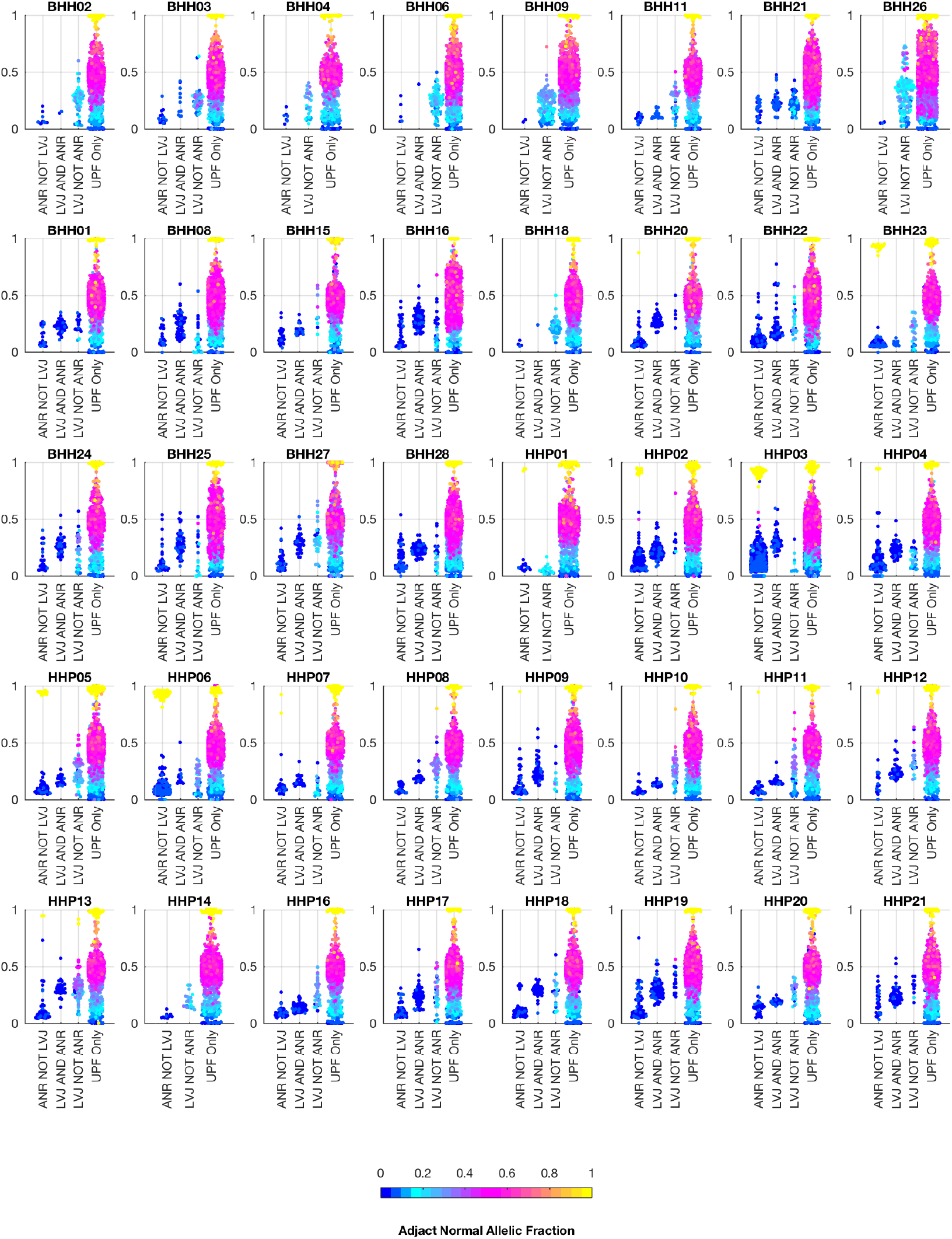
Comparison of Allelic Fractions of Variants in Archival Dataset by Calling Method. For each of the breast and prostate patients, the allele fractions in the tumor sample are plotted for the variants detected in each of the three approaches. The color of each point indicates the allele fraction of the variant in the adjacent normal sample. Most of the variants detected in the adjacent normal as reference approach, but not lumosVar 2.0 joint analysis (ANR NOT LVJ), have low allele fractions in both the tumor and the adjacent normal. The variants detected by lumosVar 2.0 joint analysis, but not adjacent normal as reference approach (LVJ NOT ANR) typically have higher allele fractions in the tumor, and lower allele fractions in the adjacent normal, though lumosVar 2.0 joint analysis also detects some variants that are lower allele fraction in the tumor and higher allele fraction in the adjacent normal in a few patients such as HPP01. The variants only called in the unmatched filtering (UPF only) approach have similar allele fractions in the tumor and adjacent normal samples. The eight patients in the top row had the adjacent normal tissue macrodissected from tumor containing slides and these patients typically have more variants detected by lumosVar 2.0 joint analysis and not ANR compared to the remaining patients whose adjacent normal sample was procured from separate slides.

We also analyzed the archival tissue using two additional approaches: 1) a filtering strategy where standard somatic variant calling tools were used against an unmatched reference, and variants found in dbSNP were excluded as likely germline, referred to as the unmatched plus filtering approach (UPF), and 2) a strategy that used the tumor adjacent normal sample as the normal reference in standard somatic variant calling tools, referred to as the adjacent normal as reference approach (ANR). Using the UPF strategy, we found that most of the variants called using the filtering strategy have variant allele fractions around 50% in both the low- and high-tumor-content samples, suggesting that most are private germline variants. Using the ANR approach, we are only identifying variants with allele fractions in the adjacent normal sample that are at or very close to zero. The variants called by lumosVar 2.0 generally have higher allele fractions in the tumor samples and low allele fractions in the adjacent normal samples, as expected (Figure 6).

In order to compare the ability of the three approaches to detect likely drivers, mutations called by any of the three approaches were compared against the Cancer Hotspots database, which reports recurrent mutations in 11,119 tumor samples (Cancer Hotspots). A total of 28 hotspot mutations were called including eight mutations with *in vitro* or *in vivo* validation (level-3), two mutations detected in the Cancer Hotspots dataset that were previously reported (level-2), and eighteen mutations that were novel in the Cancer Hotspots dataset (level-1). Of the ten level-3 and level-2 mutations, all were called in the UNF approach, lumosVar 2.0 joint analysis called eight, and only six were called in the ANR approach (Table 5). The two level-2 and level-3 mutations missed by lumosVar 2.0 had low allele fractions in the tumor sample (5-6%) and were not detected in the adjacent tissue, while the four level-3 hotspots variants missed by the ANR approach had moderate allele fraction in the tumor (17-35%) and low allele fractions in the adjacent tissue (2-16%). Seventeen of the eighteen level-1 hotspots were called only in the UNF approach, and these tended to have similar allele fractions in the tumor and adjacent normal samples. These include the same APOBR mutation called in 13 patients, and the same DHRS4 mutation called in four patients (Supplemental Table 3). Putative mutations that are common within a dataset, but not know to be common in cancer, are suggestive of alignment artifacts (Teer et al., 2017). Both the UNF approach and LumosVar 2.0 detected the eighteenth level-1 hotspot which was a CDH3 truncating mutations with high allele fractions in the tumor and low in the adjacent normal.

**Table 5.** Hotspot mutation detection.

| Patient | Gene | AA Change | AD AN (AF)<sup>a</sup> | AD TM (AF)<sup>b</sup> | LVJ | ANR | CHL<sup>c</sup> | Cosmic Count |
| --- | --- | --- | --- | --- | --- | --- | --- | --- |
| BHH01 | PIK3CA | H1047R | 473,9 (0.02) | 378,201 (0.35) | Yes | LQC<sup>d</sup> | 3 | 1806 |
| HHP13 | PIK3CA | E545K | 332,0 (0) | 195,11 (0.05) | No | Yes | 3 | 332 |
| BHH06 | AKT1 | E17K | 121,23 (0.16) | 151,62 (0.29) | Yes | No | 3 | 295 |
| BHH24 | AKT1 | E17K | 123,1 (0.01) | 118,60 (0.34) | Yes | Yes | 3 | 295 |
| BHH28 | PIK3CA | H1047L | 417,0 (0) | 348,81 (0.19) | Yes | Yes | 3 | 262 |
| HHP19 | TP53 | G245S | 186,1 (0.01) | 61,76 (0.55) | Yes | Yes | 3 | 81 |
| BHH18 | PIK3CA | Q546E | 201,16 (0.07) | 164,38 (0.19) | Yes | No | 3 | 3 |
| BHH18 | PIK3CA | G106R | 127,8 (0.06) | 96,19 (0.17) | Yes | No | 3 | 2 |
| BHH25 | PIK3CA | E726K | 269,0 (0) | 268,17 (0.06) | No | Yes | 2 | 31 |
| BHH25 | SF3B1 | K666E | 210,0 (0) | 122,46 (0.27) | Yes | Yes | 2 | 19 |

<sup>a</sup>Allelic depth of reads supporting the reference, alternate alleles in the adjacent normal sample and <sup>b</sup>tumor sample. <sup>c</sup>Cancer hotspots database validation level (Cancer Hotspots).
<sup>d</sup>Called by one of three paired somatic variant callers (strelka).

## 4 Discussion

Detecting somatic mutations when a normal tissue sample is not available remains a challenging problem. We present a method that leverages tumor-adjacent normal tissue, and is robust to significant levels of tumor contamination. Simulation studies suggest that a multi-sample approach should be more powerful than a single-sample approach, even if there is a small difference in tumor content between the two samples. Evaluation of a set of GBM samples with low tumor content (from NE biopsies) and high tumor content (from CE resections) further demonstrates the sensitivity and precision of the joint approach. Practical application of this approach to a set of FFPE breast and prostate samples shows the feasibility of this approach with typical archival samples.

Although the presence of genomic variants can be used to guide personalized medicine, not all patients with a targetable aberration for a given therapy will respond equally well to the therapy. Many factors contribute to patient response, including the presence of other aberrations affecting the same pathway, aberrations affecting alternative pathways, and sub-clonal resistance mutations. Disentangling how the interaction of genomic aberrations predicts response to a given therapy will require large sets of patients with known treatment outcomes. Banks of archival samples show great potential to accelerate such research, as medium- and long-term outcomes may already be known. The approach outlined here should enable researchers to use archival samples more effectively to answer such critical questions.

A complex relationship exists between tumor heterogeneity and clinical outcome, with moderately heterogeneous tumors having worse outcomes than both more homogenous tumors and more heterogeneous tumors (Andor et al., 2015, 2017). Measuring the overall level of heterogeneity, in addition to detecting the clonal prevalence of individual variants, can provide insight into susceptibility and resistance to targeted therapies (Saunders et al., 2012b). LumosVar 2.0’s ability to jointly analyze multiple samples from the same patient and integrate copy number and mutation data should be useful even when a matched normal sample is available to track mutations across longitudinal sample collections and spatially diverse samples, to gain insight into the tumor’s evolution. Future work will further evaluate and benchmark lumosVar 2.0’s clonal variant group analysis.

Compared to the single sample lumosVar analysis, the joint approach requires lower total sequencing coverage to obtain the same sensitivity. Based on the simulation studies, we find that if the adjacent normal tissue has less than 25% tumor cell contamination, and the tumor sample has at least 55% percent tumor cells, then only 200X total coverage (100X for each sample) is required to detect 80% of the somatic variants that are in all of the tumor cells. However, higher coverage would be desirable in order to detect low abundance sub-clonal variants. Due to the large difference in prior probabilities of homozygous reference versus somatic variants in this model, lumosVar 2.0 tends to be less sensitive to low abundance variants compared to other somatic variant callers. LumosVar 2.0 also has more stringent quality filtering than most paired somatic variant callers because the same artifacts often appear in the tumor and germline sample, so paired callers can use the presence in the germline to eliminate those artifacts. The probability that a variant is somatic or germline is calculated assuming that the allele specific copy number of the position is known with certainty, while there is clearly uncertainty in both setting the segmentation boundaries assigning both the copy number state of a given segment. Inspection of incorrectly classified variants suggests that segmentation boundary placement is a major source of error. We believe that a more sophisticated segmentation algorithm that is able to capture the uncertainty of segmentation boundary placement would yield the largest improvements in performance. We also recognize that an underlying assumption of our copy number model, that at most one copy number altered state may occur in a given segment across the patient samples, is an oversimplification, and a more realistic copy number model may improve both the copy number and variant calling results.

Though it may seem surprising that somatic variants were detected in histologically normal tumor adjacent tissue, previous studies have identified DNA, epigenetic, and gene expression alterations in tumor adjacent tissue (Troester et al., 2016). The theory of field cancerization proposes that epigenetic changes in the adjacent tissue creates a permissive environment for malignant transformation and sometimes can lead to multifocal disease and/or clonally independent recurrence. The sequencing of DNA from tumor adjacent tissue could serve a dual purpose in helping to identify somatic mutations when another source of normal tissue is not available, as well as helping to better understand the phenomenon of field cancerization.

## 5 Author Contributions

RH designed and implemented the LumosVar 2.0 software. SK designed and implemented the custom pileup engine. RH, ET, JA, WL, NH, JN, CK, RK, MB, and SB were involved in designing and generating data for the breast and prostate study. RH, NT, MB, and SB were involved in the analysis and interpretation of the GBM study. DE assisted with data analysis. SK aided in mathematical formulations. RH, SK, and SB drafted and edited the manuscript. All authors have read and approved the manuscript.

## 6 Conflict of Interest Statement

The authors declare that the research was conducted in the absence of any commercial or financial relationships that could be construed as a potential conflict of interest.

## 7 Acknowledgements

We would like to thank Nicholas Schork, David Craig, Jessica Aldrich, Austin Christofferson, and Jonathan Keats for helpful discussion. We also thank the Ben and Catherine Ivy Foundation and GE Global Research for funding for this study.

## References

Allen, E. M. V., Wagle, N., Stojanov, P., Perrin, D. L., Cibulskis, K., Marlow, S., et al. (2014). Whole-exome sequencing and clinical interpretation of FFPE tumor samples to guide precision cancer medicine. Nat. Med. 20, 682–688. doi: 10.1038/nm.3559.

Andor, N., Graham, T. A., Jansen, M., Xia, L. C., Aktipis, C. A., Petritsch, C., et al. (2015). Pan-cancer analysis of the extent and consequences of intratumor heterogeneity. Nat. Med. 22, 105–113. doi: 10.1038/nm.3984.

Andor, N., Maley, C. C., and Ji, H. P. (2017). Genomic Instability in Cancer: Teetering on the Limit of Tolerance. Cancer Res. 77, 2179–2185. doi: 10.1158/0008-5472.CAN-16-1553.

Byron, S. A., Tran, N. L., Halperin, R. F., Phillips, J. J., Kuhn, J. G., Groot, J. F. de, et al. (2017). Prospective feasibility trial for genomics-informed treatment in recurrent and progressive glioblastoma. Clin. Cancer Res., clincanres.0963.2017. doi: 10.1158/1078-0432.CCR-17-0963.

Cancer Hotspots Available at: http://cancerhotspots.org/#/home [Accessed December 15, 2017].

Cheng, D. T., Mitchell, T. N., Zehir, A., Shah, R. H., Benayed, R., Syed, A., et al. (2015). Memorial Sloan Kettering-Integrated Mutation Profiling of Actionable Cancer Targets (MSK-IMPACT): A Hybridization Capture-Based Next-Generation Sequencing Clinical Assay for Solid Tumor Molecular Oncology. J. Mol. Diagn. 17, 251–264. doi: 10.1016/j.jmoldx.2014.12.006.

Christoforides, A., Carpten, J. D., Weiss, G. J., Demeure, M. J., Hoff, D. D. V., and Craig, D. W. (2013). Identification of somatic mutations in cancer through Bayesian-based analysis of sequenced genome pairs. BMC Genomics 14, 302. doi: 10.1186/1471-2164-14-302.

Cibulskis, K., Lawrence, M. S., Carter, S. L., Sivachenko, A., Jaffe, D., Sougnez, C., et al. (2013). Sensitive detection of somatic point mutations in impure and heterogeneous cancer samples. Nat. Biotechnol. 31, 213–219. doi: 10.1038/nbt.2514.

DePristo, M. A., Banks, E., Poplin, R., Garimella, K. V., Maguire, J. R., Hartl, C., et al. (2011). A framework for variation discovery and genotyping using next-generation DNA sequencing data. Nat. Genet. 43, 491–498. doi: 10.1038/ng.806.

Dotto, G. P. (2014). Multifocal epithelial tumors and field cancerization: stroma as a primary determinant. J. Clin. Invest. 124, 1446–1453. doi: 10.1172/JCI72589.

Frampton, G. M., Fichtenholtz, A., Otto, G. A., Wang, K., Downing, S. R., He, J., et al. (2013). Development and validation of a clinical cancer genomic profiling test based on massively parallel DNA sequencing. Nat. Biotechnol. 31, 1023–1031. doi: 10.1038/nbt.2696.

Halperin, R. F., Carpten, J. D., Manojlovic, Z., Aldrich, J., Keats, J., Byron, S., et al. (2017). A method to reduce ancestry related germline false positives in tumor only somatic variant calling. BMC Med. Genomics 10, 61. doi: 10.1186/s12920-017-0296-8.

Hanahan, D., and Weinberg, R. A. (2000). The Hallmarks of Cancer. Cell 100, 57–70. doi: 10.1016/S0092-8674 (00)81683-9.

Heaphy, C. M., Griffith, J. K., and Bisoffi, M. (2009). Mammary field cancerization: molecular evidence and clinical importance. Breast Cancer Res. Treat. 118, 229–239. doi: 10.1007/s10549-009-0504-0.

Hoang, M. L., Kinde, I., Tomasetti, C., McMahon, K. W., Rosenquist, T. A., Grollman, A. P., et al. (2016). Genome-wide quantification of rare somatic mutations in normal human tissues using massively parallel sequencing. Proc. Natl. Acad. Sci. U. S. A. 113, 9846–9851. doi: 10.1073/pnas.1607794113.

Jones, S., Anagnostou, V., Lytle, K., Parpart-Li, S., Nesselbush, M., Riley, D. R., et al. (2015). Personalized genomic analyses for cancer mutation discovery and interpretation. Sci. Transl. Med. 7, 283ra53. doi: 10.1126/scitranslmed.aaa7161.

Kalatskaya, I., Trinh, Q. M., Spears, M., McPherson, J. D., Bartlett, J. M. S., and Stein, L. (2017). ISOWN: accurate somatic mutation identification in the absence of normal tissue controls. Genome Med. 9, 59. doi: 10.1186/s13073-017-0446-9.

Khurana, E., Fu, Y., Chakravarty, D., Demichelis, F., Rubin, M. A., and Gerstein, M. (2016). Role of non-coding sequence variants in cancer. Nat. Rev. Genet. 17, 93–108. doi: 10.1038/nrg.2015.17.

Marrone, M., Schilsky, R. L., Liu, G., Khoury, M. J., and Freedman, A. N. (2015). Opportunities for Translational Epidemiology: The Important Role of Observational Studies to Advance Precision Oncology. Cancer Epidemiol. Prev. Biomark. 24, 484–489. doi: 10.1158/1055-9965.EPI-14-1086.

Martincorena, I., Roshan, A., Gerstung, M., Ellis, P., Loo, P. V., McLaren, S., et al. (2015). High burden and pervasive positive selection of somatic mutations in normal human skin. Science 348, 880–886. doi: 10.1126/science.aaa6806.

Nonn, L., Ananthanarayanan, V., and Gann, P. H. (2009). Evidence for Field Cancerization of the Prostate. The Prostate 69, 1470–1479. doi: 10.1002/pros.20983.

Saunders, C. T., Wong, W. S. W., Swamy, S., Becq, J., Murray, L. J., and Cheetham, R. K. (2012a). Strelka: accurate somatic small-variant calling from sequenced tumor–normal sample pairs. Bioinformatics 28, 1811–1817. doi: 10.1093/bioinformatics/bts271.

Saunders, N. A., Simpson, F., Thompson, E. W., Hill, M. M., Endo-Munoz, L., Leggatt, G., et al. (2012b). Role of intratumoural heterogeneity in cancer drug resistance: molecular and clinical perspectives. EMBO Mol. Med. 4, 675–684. doi: 10.1002/emmm.201101131.

Sherry, S. T., Ward, M. H., Kholodov, M., Baker, J., Phan, L., Smigielski, E. M., et al. (2001). dbSNP: the NCBI database of genetic variation. Nucleic Acids Res. 29, 308–311.

Teer, J. K., Zhang, Y., Chen, L., Welsh, E. A., Cress, W. D., Eschrich, S. A., et al. (2017). Evaluating somatic tumor mutation detection without matched normal samples. Hum. Genomics 11, 22. doi: 10.1186/s40246-017-0118-2.

Troester, M. A., Hoadley, K. A., D’Arcy, M., Cherniack, A. D., Stewart, C., Koboldt, D. C., et al. (2016). DNA defects, epigenetics, and gene expression in cancer-adjacent breast: a study from The Cancer Genome Atlas. NPJ Breast Cancer 2, 16007. doi: 10.1038/npjbcancer.2016.7.

Waldron, L., Ogino, S., Hoshida, Y., Shima, K., Reed, A. E. M., Simpson, P. T., et al. (2012). Expression Profiling of Archival Tumors for Long-term Health Studies. Clin. Cancer Res. 18, 6136–6146. doi: 10.1158/1078-0432.CCR-12-1915.

Wei, L., Papanicolau-Sengos, A., Liu, S., Wang, J., Conroy, J. M., Glenn, S. T., et al. (2016). Pitfalls of improperly procured adjacent non-neoplastic tissue for somatic mutation analysis using next-generation sequencing. BMC Med. Genomics 9, 64. doi: 10.1186/s12920-016-0226-1.

